# Copy number variations of *Plasmodium vivax DBP1*, *EBP/DBP2*, and *RBP2b* in Duffy-positive and Duffy-negative Ethiopians

**DOI:** 10.1101/2024.04.29.591731

**Authors:** Kareen Pestana, Anthony Ford, Rei Rama, Beka Abagero, Daniel Kepple, Junya Tomida, Jean Popovici, Delenasaw Yewhalaw, Eugenia Lo

## Abstract

Recent evidence challenges the belief that Duffy-negative individuals are resistant to *Plasmodium vivax* due to lacking Duffy Antigen Receptor for Chemokines (DARC). Erythrocyte Binding Protein (EBP/DBP2) has shown moderate binding to Duffy-negative erythrocytes *in vitro*. Reticulocyte Binding Protein 2b (RBP2b) interactions with Transferrin Receptor 1 (TfR1) suggest involvement in Duffy-negative infections. Gene copy number variations in *PvDBP1*, *PvEBP*/*DBP2*, and *PvRBP2b* were investigated in Duffy-positive and Duffy-negative *P. vivax*-infected individuals from Ethiopia. Among Duffy-positive samples, 34% displayed *PvDBP1* duplications (Cambodian-type). In Duffy-negative infections, 30% showed duplications, mostly Cambodian-type. For *PvEBP*/*DBP2* and *PvRBP2b*, Duffy-positive samples exhibited higher duplication rates (1-8 copies for *PvEBP*/*DBP2*, 1-5 copies for *PvRBP2b* 46% and 43% respectively) compared to Duffy-negatives (20.8% and 26% respectively). The range of copy number variations was lower in Duffy-negative infections. Demographic and clinical factors associated with gene multiplications in both Duffy types were explored, enhancing understanding of *P. vivax* evolution in Duffy-negative Africans.

## Introduction

Malaria presents a continuing health burden in developing countries, particularly those in sub-Saharan Africa where 94% of global malaria cases and deaths occur [1]. Majority of the people of Sub-Saharan African descent have -67T>C mutation (rs2814778) in the GATA-1 transcription factor binding site of the Duffy Antigen for Chemokines (*DARC)* gene, leading to reduced expression of the receptor on the surface of their red blood cells, and thus low susceptibility to *P. vivax* infection [2]. Nevertheless, several recent studies show that Duffy-negative individuals can be infected by *P. vivax* and symptomatic and asymptomatic *P. vivax* infections are increasingly reported across Africa [3]. It is clear that *P. vivax* is capable of infecting and causing malaria disease in Duffy-negative individuals, though these infections generally have lower parasitemia than the Duffy-positive ones [4].

The interaction between *P. vivax* Duffy Binding Protein 1 (PvDBP1) and DARC is a well-known Duffy-dependent pathway for invasion of human erythrocytes. The invasion mechanisms of *P. vivax* in Duffy-negative erythrocytes, however, are largely unclear. Previous studies showed that *P. vivax* erythrocyte binding protein (PvEBP/DBP2), a homologue of PvDBP1, shares similar binding domains and binds moderately to Duffy-negative reticulocytes [5]. The binding receptor for PvEBP/DBP2 was recently identified to be Complement Receptor 1 (CR1), presenting evidence for an alternative DARC-independent pathway of erythrocyte invasion [6]. Given *P. vivax* prefers binding to reticulocytes, *P. vivax* reticulocyte binding protein 2 (PvRBP2b) has also been shown to bind to transferrin receptor 1 (TfR1) on the surface of Duffy-positive reticulocytes and contribute to a non-Duffy invasion pathway [7].

*PvDBP1* in the African *P. vivax* had higher copy number variation than the Southeast Asian and South American isolates [8, 9]. Compared with *P. vivax* from Cambodia (1-3x copies with 37% had >1 copies), the Malagasy *P. vivax* showed a greater variation in *PvDBP1* copy number ranging from 1-5x copies with 45% of the isolates had high-order copies [8]. In Ethiopia and Madagascar, Duffy-positive and Duffy-negative individuals co-exist within the same area. Among the Ethiopian Duffy-positives, *PvDBP1* ranged from 1-2+x copies with 64% had >1 copies [10]. In *P. vivax* from Duffy-negatives, more than five copies of *PvDBP1* were detected previously, though only two samples were examined [5]. Like *PvDBP1*, higher copy number of *PvEBP/DBP2* has been reported in the Malagasy (1-5 copies with 44% had >1 copies) and Cambodian (1-2 copies with 83% had >1 copies) *P. vivax* [8], but is less characterized in Duffy-negative populations. It is yet unclear whether copy number variations occur in *PvRBP*2b of the African *P. vivax*.

Copy number variations have been reported in a multitude of genes in other *Plasmodium* species with different effects on antimalarial resistance and immune evasion. Gene multiplications appear to occur non-randomly and distributed throughout the *Plasmodium* genome. In *P. falciparum*, high copies of the multidrug resistance gene 1 (*pfmdr1*) was associated with resistance to mefloquine, halofantrine, and quinine [11–13]. Multiple copies of *var* genes that encode antigens from the intra-erythrocytic stage of *P. falciparum* were shown with greater levels of nucleotide diversity, allowing for greater immune evasion responses [14, 15]. Further, mitochondrial genes (*cytochrome b*), apicoplast genes (*clcp*), erythrocyte surface proteins (*rifin* gene families), erythrocyte binding proteins (*stevor* gene families), and folate pathway enzymes (*gch1*) of *P. falciparum* that may relate to parasite development and host invasion were detected with multiple copies [16–18]. These variations have clinical implications in both treatment of the disease and development of vaccine targets. In *P. vivax,* similar patterns of gene multiplications have been reported. Like *pfmdr1*, higher copies of *P. vivax mdr*1 (*pvmdr1*) were shown to correlate with mefloquine resistance [19–21]; and high copies of *PvDBP1* offer protection to parasite growth *in vitro* against anti-PvDBP antibodies [22]. Among several antigens tested for humoral responses, *PvEBP/DBP2* showed a greater and more long-lasting response than *PvDBP1* [23]. Other genes such as the 28S ribosomal RNA gene and a gene that encodes an exported *Plasmodium* proteins also showed copy number variations in the Ethiopian *P. vivax* [24].

The frequency of *PvDBP1* and *PvEBP/DBP2* copy number variations (CNV) in Duffy-negative infections remains unclear. It is also unknown whether *PvRBP2b* shows similar trends of CNV in Duffy-positive and Duffy-negative individuals. This study aims to 1) investigate CNV of *PvDBP1*, *PvEBP/DBP2*, and *PvRBP2b* in the Ethiopian *P. vivax* isolates; 2) compare CNV of these three genes between Duffy-positive and Duffy-negative individuals; and 3) determine the association of *PvDBP1*, *PvEBP/PvDBP2*, and *RBP2b* CNV with parasitemia, transcriptomic levels, and demographic and clinical features. Information on the structure of genes associated with host erythrocytic invasion sheds light on potentially novel invasion pathway(s) and impacts by *P. vivax* in Duffy-negative Africans.

## Materials and Methods

### Sample collection and molecular screening

Finger-prick blood was collected from 155 *P. vivax*-infected patients in Ethiopia. DNA was extracted from the dried blood spot using the Saponin/Chelex method [25]. Additionally, at the time of sample collection, health and socio-demographic information including gender, age, body temperature, and malaria were recorded. Identification and estimation of *P. vivax* parasitemia was performed using the 18S rRNA SYBR green quantitative PCR method as previously described [26–28] (**Supplementary Table 1**). Each assay utilized 10-fold serial dilution of *P. vivax* Pakchong (MRA-342G) and Nicaragua (MRA-340G) isolates as positive controls, and water and non-infected samples as negative controls. Samples yielding *Ct* values higher than 40 were considered negative for *P. vivax*. Parasitemia was estimated using the following equation: Log parasite density per μl of blood = 2^Ex(40−^ *^Ct^* ^sample)^/10. For samples identified as *P. vivax* positive, qPCR-based TaqMan assay (**Supplementary Table 1**) was used to genotype SNP position rs2814778 (-67T>C) of the *DARC* gene to determine whether the patient was homozygous (T/T) or heterozygous (C/T) Duffy-positive or Duffy-negative (C/C) [29]. Samples identified as Duffy-negative (C/C) were further PCR amplified for the *DARC* genes and verified by Sanger sequencing.

### Detecting gene duplications by PCR and qPCR assays

For *PvDBP*1, the detection of the Cambodian- or Malagasy-type duplications was based on previously published PCR protocols [9, 10]. Each PCR reaction contained 10 μl of DreamTaq™ Green PCR Master Mix (2X) (Thermo Fisher Scientific, USA), 0.5 μM of each primer (**Supplementary Table 1**), and 3 μl of DNA template. PCR conditions were 94°C for 2 minutes in initial denaturation, followed by 35 cycles of 94°C for 20 seconds, 55°C for 30 seconds, 68°C for 1 minute and a final extension for 4 minutes.

To determine copy number variation of *PvDBP1*, *PvEBP/DBP2*, and *PvRBP2b,* a qPCR-based protocol was optimized based on previously established protocols for each gene using plasmids as standard curve controls. Each sample was normalized using *P. vivax* β*-tubulin* as an endogenous control (a single-copy gene as a reference standard) [8, 30]. β*-tubulin* allowed for relative quantification of *PvDBP1*, *PvEBP/DBP2*, and *PvRBP2b* in each sample without requiring prior normalization of the amount of parasite DNA in a sample. Plasmids that contain β*-tubulin* (cloned in pUC57)*, PvDBP1* (in pUC57)*, PvEBP/DBP2* (in pUC57), and *PvRBP2b* (cloned in a pEX vector) were synthesized by GenScript Biotech. We used different proportions of plasmid mixes based on previous studies to capture a large range of copy number variations in African samples [22]. The plasmid mixes were prepared with a 1:1 ratio of β*-tubulin* and the gene of interest (i.e. 50µL of 0.1 pg/µL of β*-tubulin* and 50µL of 0.1 pg/µL of *PvDBP* to represent 1 copy of the reference gene to 1 copy of the gene of interest). The proportion of gene of interest to β*-tubulin* increased to represent higher copy numbers (i.e. 25µL of 0.1 pg/µL of β*-tubulin* and 75µL of 0.1 pg/µL of *PvDBP1* plasmid to represent 1 copy of β*-tubulin* per every 3 copies of *PvDBP1*) [8]. The assays for *PvDBP1* and *PvEBP*/*DBP2* have been previously verified by whole genome sequencing for copy number validation, although whole genome sequencing for copy number variation for PvRBP2b is still underway [8].

The amplification was carried out in 20 μl reactions containing 10 μl of PowerUp™ SYBR Green Master Mix (Thermo Fisher, US), 0.5 μM of each primer (**Supplementary Table 1**) and 3 μl of DNA template. The PCR conditions were 95°C for 15 minutes followed by 45 cycles of 95°C for 15 seconds, 60°C for 20 seconds, 72°C for 20 seconds and a melt curve stage on a QuantStudio 3 quantitative PCR system (Thermo Fisher) and each assay was performed in triplicate. Relative quantification was performed using a range of 1:1 to 1:11 ratio of β-tubulin and plasmid containing the gene of interest to calculate a standard curve (Thermo Fisher Design and Analysis software). Expression analysis calculation was performed using the ΔΔCq method, ΔC_T_ = C_T_ of target gene to C_T_ of control gene (β*-tubulin)*.

### Association analyses with socio-demographic, clinical, and gene expression data

Individuals infected with *P. vivax* potentially experienced a wide array of symptoms, which may be a function of parasitemia levels, Duffy blood group, and duplications in *PvDBP*1, *PvEBP/DBP2*, and *PvRBP2b*. These symptoms include, but not limited to, chills and shivering, fatigue, malaise, muscle pain, joint pain, headache, irritability, nausea, vomiting and loss of appetite. Simple logistic regressions were used to obtain the adjusted odds ratio to determine whether copy number variation was correlated to socio-demographic factors, clinical symptoms, and gene expression data, respectively, in Duffy positive and Duffy negative individuals. To predict the likelihood of *P. vivax* infected individual experiencing any of the clinical symptoms, we constructed a multivariate logistic regression model as a function of log parasitemia, age, sex, Duffy type (T/T or T/C Duffy-positive and C/C Duffy-negative), and gene copies observed in *PvDBP*1, *PvEBP/DBP2,* and *PvRBP2b* for each symptom using SAS v9.4 giving a total of 10 classifiers. In each model, clinical symptom for each subject was coded as 1 (present) or 0 (absent of symptom). For categorical predictor variables, we used dummy coding rather than the default effects coding in SAS. Because of limited sample size, some models did not include the categorical predictors DuffyType and/or gender. We calculated the odds ratios, along with corresponding 95% confidence intervals of whether a patient experienced the clinical symptom for each predictor variable, regardless of statistical significance. We conducted transcriptomic analysis using the identical sample set from a preceding study, comparing gene expression profiles with estimates of copy number variation to determine whether copy number variation had an effect on gene expression profiles of invasion genes [31].

## Results

### Copy number variations of *PvDBP1*, *PvEBP*/*DBP2*, and *PvRBP2b*

Of 155 *P. vivax* confirmed samples, 109 were Duffy-positives (55 T/T and 54 T/C) and 46 were Duffy-negatives. In general, 45-53% of Duffy-positives with *P. vivax* infections had multi-copy for *PvDBP1*, *PvEBP/DBP2*, and *PvRBP2b*, whereas only ∼25% of Duffy-negatives with *P. vivax* infections had multi-copy for these genes (**Table 1**; **Supplementary Table 2**). For *PvDBP1*, 47.3% (26/55) T/T homozygous and 53.7% (29/54) T/C heterozygous Duffy-positive individuals had multiple copies. All of them were detected with Cambodian-type duplications. By contrast, 34.8% (16/46) of Duffy-negative individuals had multiple copies. The majority of these multi-*DBP1* samples (15/16) had the Cambodian-type duplications except one that was detected with the Malagasy-type duplications. *PvDBP1* ranged from 1-11 copies in T/T homozygous Duffy-positives, 1-5 copies in T/C heterozygous Duffy-positives, and up to three copies in Duffy-negatives (**Figure 1**). The number of *PvDBP1* copies were significantly higher in T/T homozygous than T/C heterozygous Duffy-positive (*p*=0.0115) and C/C homozygous Duffy-negative individuals (*p*<0.001; **Figure 1**). No significant difference was detected in parasitemia between parasites of single-copy and multi-copy *PvDBP1* in both Duffy-positive (T/T and T/C; *p*=0.62 and 0.92, respectively) and Duffy-negative (C/C; *p*=0.78) infected individuals (**Figure 2**).

**Figure 1:**
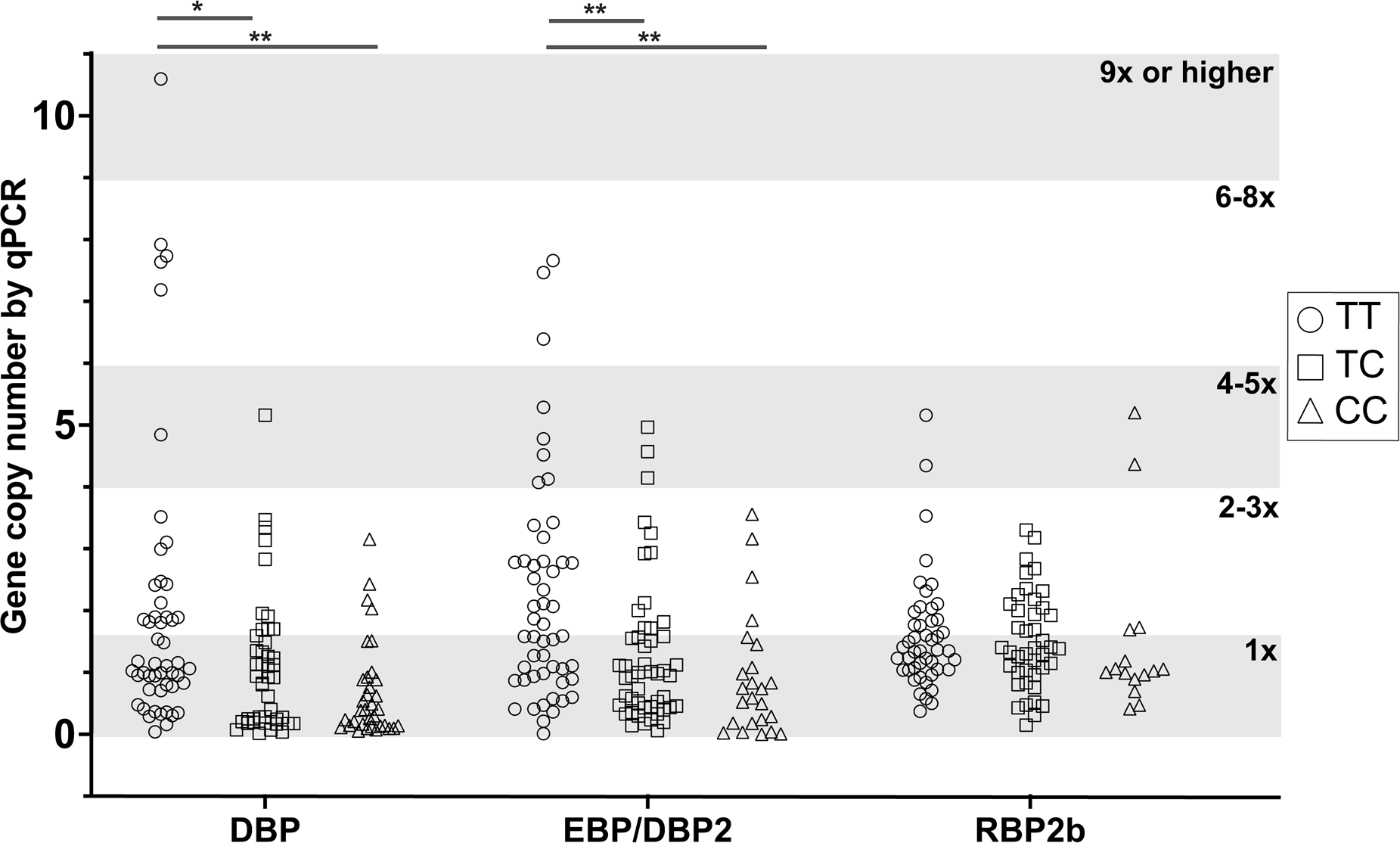
Copy number variation of *PvDBP1*, *PvEBP*/*DBP2*, and *PvRBP2b* between Duffy-positive (T/T and T/C) and Duffy-negative (C/C groups). The significance levels are denoted by asterisks, with ‘*’ representing p-values where *P* ≤ 0.05 and ‘**’ for *P* ≤ 0.01. These findings underscore the impact of Duffy Genotypes on gene copy number variations, particularly accentuated in the case of *PvDBP* and *PvEBP*/*DBP2* genes.

**Figure 2:**
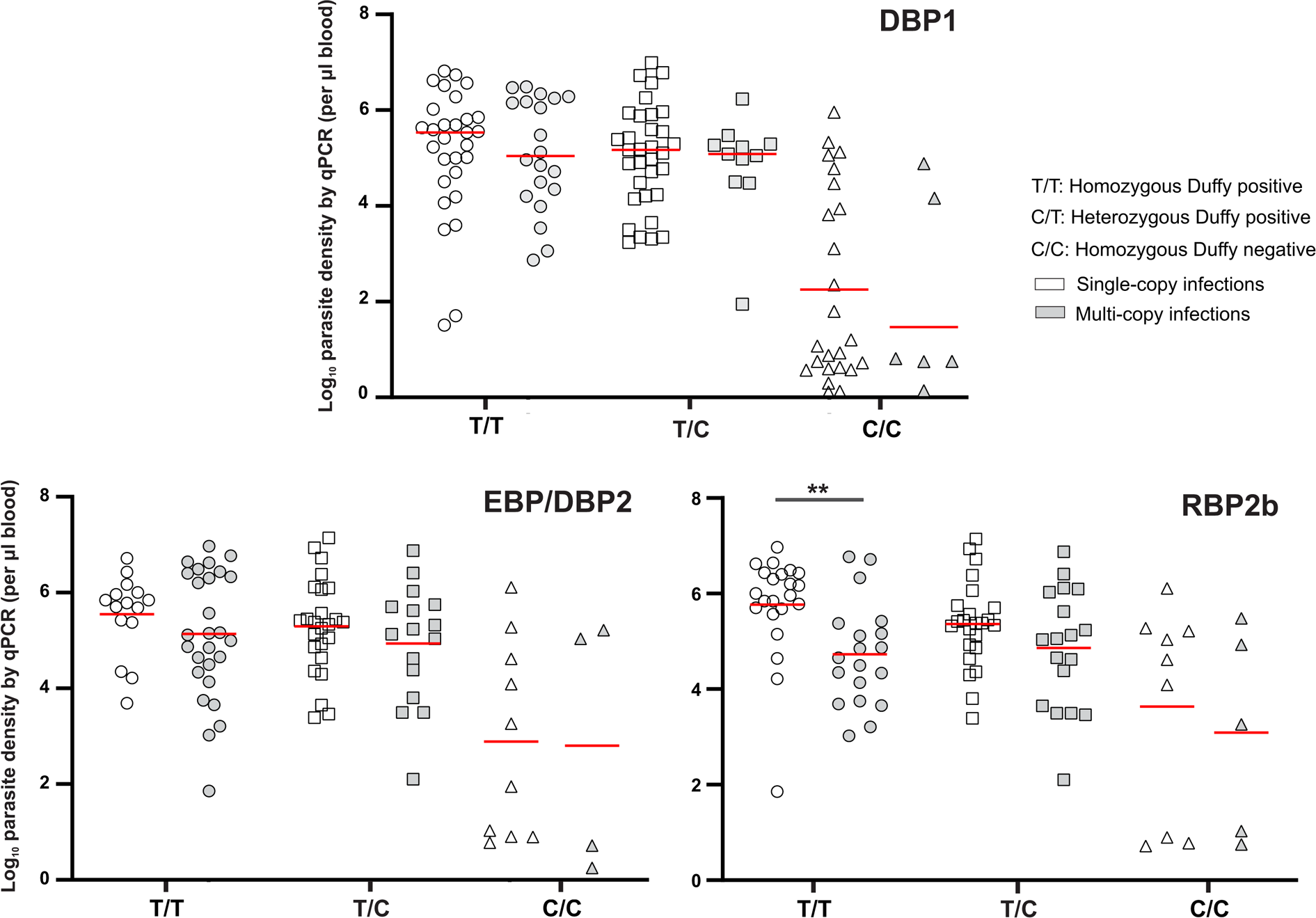
Comparison of single-copy versus multi-copy *PvDBP1*, *PvEBP*/*DBP2*, and *PvRBP2b* and estimated log parasitemia for Duffy-positive (T/T and T/C) and Duffy-negative (C/C) groups. Duffy-negative groups regardless of gene multiplicity had lower parasitemia than Duffy-positive groups. Significant differences in parasitemia between single-copy and multi-copy PvDBP1 parasites were not detected across both Duffy-positive groups (T/T and T/C; p=0.62 and 0.92, respectively) and Duffy-negative individuals (C/C; p=0.78). Similarly, no significant distinctions in parasitemia were observed between single-copy and multi-copy infections for all Duffy groups with PvEBP/DBP2 (p>0.05). However, a notable exception emerged in the T/T homozygous Duffy-positive subgroup, where a significant parasitemia difference was evident between single-copy and multi-copy infections (p=0.001).

For *PvEBP/DBP2*, 58.82% (30/51) of the T/T homozygous and 32.65% (16/49) T/C heterozygous Duffy-positive individuals had multiple copies. By contrast, 20.8% (5/24) of the Duffy-negative individuals had multiple copies (**Table 1**). *PvEBP/DBP2* copy number ranged from 1-8 copies in most samples, average 2-3 copies among T/T homozygous Duffy-positives, and up to 4-5x copies in T/C heterozygous Duffy-positives and Duffy-negatives (**Figure 1**). There were significant differences in *PvEBP/DBP2* copy number between T/T homozygous Duffy-positives and C/C homozygous Duffy-negative individuals (*p*<0.001) and T/T homozygous and T/C heterozygous Duffy-positives (*p*=0.003) (**Figure 1**). No significant difference was observed in parasitemia between the single-copy or multi-copy infections for all Duffy groups (*p*>0.05) (**Figure 2**).

For *PvRBP2b*, 43.5% (20/46) of the T/T homozygous and 42.9% (18/42) T/C heterozygous Duffy-positive individuals had multiple copies. By contrast, a lower proportion (4/15; 26.7%) of Duffy-negative individuals had multiple copies (**Table 1**). *PvRBP*2b copy number generally ranged from 1-5 copies, with up to 4-5x copies in both Duffy-positive and up to 5 copies in Duffy-negative individuals. There were no significant differences in *PvRBP2b* copy number among Duffy groups (**Figure 1**). Nevertheless, there was a significant difference in parasitemia between single-copy and multi-copy infections for the T/T homozygous Duffy positive individuals (*p*=0.001; **Figure 2**).

### Association with clinical and host factors

No significant difference was found in *PvDBP1* and *PvEBP*/*DBP2* copy number among infections of different age groups and gender (**Figure 3**). For *PvRBP2b*, a significant difference was detected in gene copy number between individuals aged 5-13 and >14 (*p*=0.025), despite small sample size was for individuals under 5 years old for all three genes.

**Figure 3:**
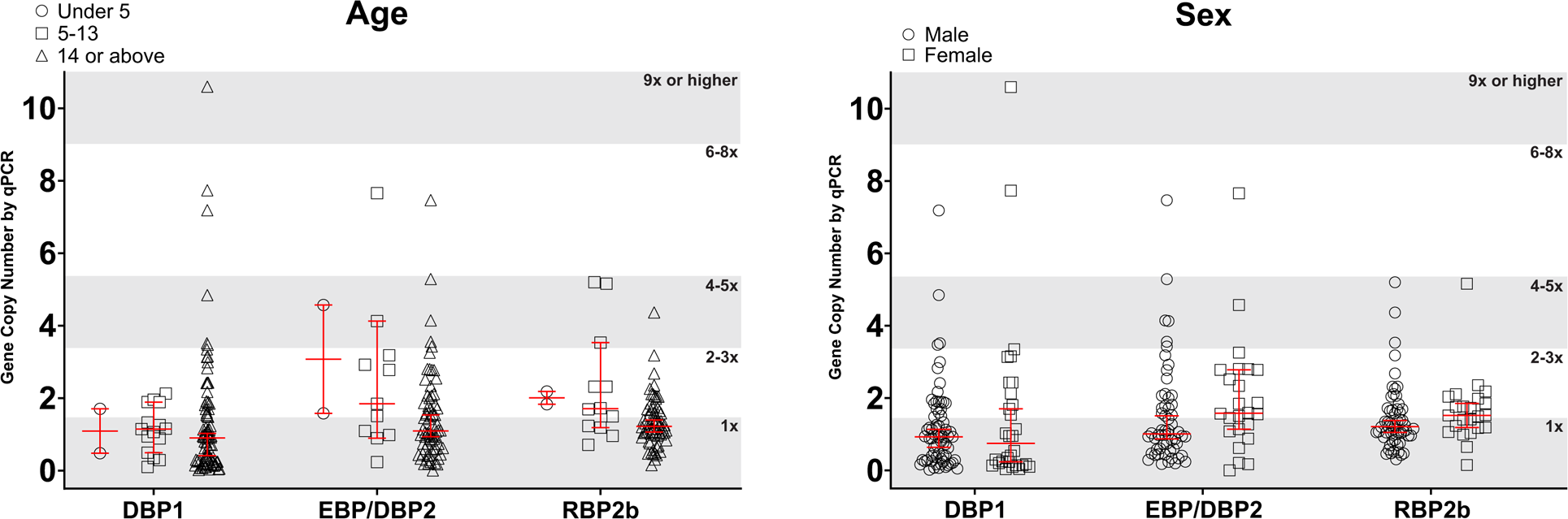
Estimated copy number variation of *PvDBP1*, *PvEBP*/*DBP2*, and *PvRBP2b* by age group. The age groups are as follows for *PvDBP1*: under 5 (*N* = 2), 5-13 (*N* = 16), above 14 (*N* = 82); *PvEBP*/*DBP2*: under 5 (*N* = 2), 5-13 (*N* = 12), above 14 (*N* = 67); and *PvRBP2b*: under 5 (*N*=2), 5-14 (*N* = 13), above 14 (*N* = 65). Figure 3b. Estimated copy number variation of *PvDBP1*, *PvEBP*/*DBP2*, and *PvRBP2b* by gender. The distribution of gender by gene copy number variation is as follows for *PvDBP1*: Female (*N* = 30) and Male (*N* = 69); *PvEBP*/*DBP2*: Female (*N* = 24) and Male (*N* = 57); and *PvRBP2b*: Female (*N* = 23) and Male (*N* = 57).

Although not significant, there was association for *P. vivax*-infected patients to experience chills and shivering symptoms when parasites had higher *PvDBP1* copies (OR of 1.41; *p*=0.07, 95% CI:0.97-2.03; **Table 2**). Similarly, patients with higher *PvEBP/DBP2* copies were 1.7-fold (*p*=0.067, 95% CI:0.96-3.01) more likely to experience muscle pain. Conversely, the odd ratio for muscle pain was low for those with *PvRBP*2b multiplications (OR:0.29; *p*=0.051, 95% CI:0.08-1.01). We found significant associations for *P. vivax*-infected patients who had *PvEBP/DBP2* (OR:0.41, *p*=0.005, 95% CI:0.22-0.77) and *PvRBP*2b multiplications (OR:3.5, *p*=0.017, 95% CI:1.25-9.75) to experience joint pain (**Table 2**).

Multiple logistic regression analyses results showed no significant association between clinical symptoms and variables including parasitemia, Duffy blood type, age, gender, and *PvDBP1*, *PvEBP/DBP2*, and *PvRBP2b* copy number in *P. vivax*-infected patients. There were associations between nausea and host factors including age (OR:1.06, *p*=0.047, 95% CI:1.00-1.12) and Duffy blood type (TT: OR of 0.037, *p*=0.009, 95% CI:0.003-0.44; and T/C: OR of 0.031, *p*=0.007, 95% CI:0.002-0.38) (**Table 2**).

### Gene copy number and expression

For a subset of 10 samples, we obtained transcriptomic data for *PvDBP1*, *PvEBP*/*DBP2*, and *PvRBP2b* to compare expression level with gene copy number. The copy number variations did not show a significant correlation with their respective expression level i.e., the TPM reads during the majority trophozoite and schizont stages (**Supplemental Table 3)**. Although *PvDBP1* and *PvRBP2b* were all single copy among the infections, transcripts per million (TPM) varied greatly, ranging from 89-324 TPM for *PvDBP1* and 22-119 TPM for *PvRBP2b* **(Supplementary Table 3)**. While several samples had two copies of *PvEBP/DBP2*, there was no significant correlation with gene expression level **(Supplementary Table 3)**.

## Discussion

The critical role of the PvDBP1-DARC interaction in Duffy-positive erythrocyte invasion by *P. vivax* has previously been documented, but the exact invasion pathway of Duffy-negative erythrocytes has yet to be elucidated [5, 9]. One hypothesis is that the multiplication of PvDBP1 could enable *P. vivax* to identify and attach to Duffy-negative erythrocytes exhibiting weak DARC expression or possibly to an unknown receptor with lower affinity to DBP1 compared to DARC. This was supported by the higher *PvDBP1* copies observed in *P. vivax* from the Malagasy Africans [7, 9], although the functional significance of *PvDBP1* multiplications remains to be investigated.

In Ethiopia, *P. vivax* from both Duffy-positive and Duffy-negative individuals had single and multi-copy *PvDBP1*, *PvEBP/DBP2*, and *PvRBP2b*. Our findings confirm the presence of Cambodian- and Malagasy-type *PvDBP1* duplications in Ethiopian Duffy-positive and Duffy-negative individuals [9], though the most prevalent duplication type was the Cambodian duplication. The Malagasy duplication was present in a single Duffy-negative patient. Different geographic regions in Ethiopia may have different prevalence of Cambodian and Malagasy duplications due variations in transmission intensity. A greater range of *PvDBP1* duplications was observed in Duffy-positive than Duffy-negative groups, although an increase in *PvDBP1* copies was not associated with increased parasitemia. It is possible that *PvDBP1* duplications offer another functional advantage such as host immune evasion. The previous hypothesis that additional multiplications of *PvDBP1* may result in greater invasion capabilities of Duffy-negative erythrocytes does not appear to be true. Additionally, dependence on 18S qPCR data obtained from blood samples for estimating parasitemia may predominantly capture the erythrocytic stage infection, potentially leading to an incomplete representation of total parasitemia. Further investigation of expanded samples in broad ecological and transmission regions of Africa is needed to test if the Malagasy duplication is more common among Duffy-negatives and/or areas of high transmission. More transcriptomic data needs to be obtained in both single-copy and mutli-copy *DBP1* infections to determine whether a change in copy number is associated with gene expression levels. Additionally, the constraints inherent in estimating copy number variation through qPCR include its potential inability to discern polyclonal infections, resulting in data that represents an average of copy number variation across different clones.

Compared to *P. vivax* from Cambodia (1-2 copies) and Madagascar (1-5 copies), the range of *PvEBP/DBP2* CNVs in the Ethiopian isolates was much higher (1-8 copies) [7]. Our findings further showed that Duffy-positive individuals had a greater prevalence and range of *PvEBP/DBP2* duplications than the Duffy-negative ones. Previous studies showed that different Duffy-positive phenotypes had differential levels of *P. vivax* incidence with those positive for the A allele (FY*A/*A) associated with a higher incidence [31[32]; such variations in host susceptibility could be related to differences in *PvDBP1* and *PvEBP/DBP2* copies. Additionally, functional studies have shown that *PvEBP/DBP2* can bind strongly to Duffy-positive reticulocytes and moderately to Duffy-negative erythrocytes [5, 33]. It remains unclear whether higher *PvEBP/DBP2* copies confers stronger binding affinity for the parasites.

This is the first study to characterize *PvRBP2b* gene amplification in Duffy-positive and Duffy-negative Africans. As *PvRBP2b* interacts with TfR1 receptors and is not dependent on interactions with DARC receptors [6], *PvRBP2b* could play a critical role in Duffy-negative erythrocyte invasion. Similar to *PvDBP1* and *PvEBP/DBP2*, *PvRBP2b* gene multiplications were more common and showed higher copies in Duffy-positive than Duffy-negative *P. vivax*. However, it is noteworthy that such differences may be attributed to small samples of Duffy-negatives, especially those with very low parasitemia that we could not amplify accurately. We are currently developing a highly sensitive digital PCR assay to precisely detect copy numbers in low-parasitemia samples and overcome this challenge.

Despite small sample size, higher *PvEBP/DBP2* copies did not show increased gene expression level, suggesting that increase in gene dosage may not necessarily increase transcription or protein production. This could be explained by first, epigenetic regulatory mechanisms that control transcription and protein synthesis, and that increased gene dosage does not impact transcription. Previous studies in *P. falciparum PfRH*4 indicated a hierarchical relationship between different erythrocytic invasion pathways and that epigenetic switching can alter which pathway is utilized, and in turn increase the parasite adaptability to host erythrocyte receptors [34]. Second, feedback between host and parasite may also influence gene expression independently from gene duplication [35, 36]. For instance, *PfEMP*1, encoded by ∼60 *var* genes, can be differentially expressed and allow for antigenic variation and host immune evasion. The positive and negative feedbacks between parasite and host could control *PfEMP*1 expression and contribute to differences between gene expression among individuals. Third, gene duplications may not correlate with transcriptomic or proteomic data if there are post-transcriptional regulation pathways that alter mRNA stability or microRNAs. In *P. falciparum,* many RNA-Binding proteins have increased expression during the intraerythrocytic life cycle, suggesting a vital role of RNA-Binding proteins in determining mRNA stability [37]. Although *Plasmodium* has been shown to lack microRNAs and RNA interference machinery [38], there is some evidence that gene expression in *Plasmodium* may be modulated by human and *Anopheles* microRNAs [39–41].

The lack of differences in parasitemia between single-copy and multi-copy infections for *PvDBP1*, *PvEBP*/*DBP2*, or *PvRBP2b* regardless of Duffy status may suggest that copy number variation of these invasion genes may not increase erythrocyte invasion or parasite affinity to host receptors. Gene duplication has been shown to serve multiple purposes in Apicomplexan parasites. *Plasmodium falciparum var* genes multiply to generate antigenic diversity and duplicated segments can recombine to generate ‘duplication chimeras’ and antigenic structures [42]. Gene duplication and diversification in the *var* gene family and the rhoptry-associated protein complex may also allow polymorphisms and/or deleterious mutations to accumulate in duplicated copies of a gene while retaining similar or overlapping functions [43, 44]. Increase in gene copy number variation has been shown to associate with disease severity. In *Toxoplasma gondii*, rhoptry protein 5 has 4-6 copies, resulting in various interactions between ROP5 and host cells that lead to differences in virulence [45]. The *Trypanosoma cruzi* genome contains many gene families including the trans-sialidase family with high copy number variation and is correlated with pathogen virulence [46, 47]. Increased copy number of *pfmdr1* showed increased resistance to mefloquine in *P. falciparum* malaria [12]. While gene duplication has the possibility of increasing parasite fitness this can come at a cost. The average parasitemia of single-copy infections was slightly higher in *PvDBP1* and *PvEBP*/*DBP2* and statistical significance was attained exclusively for *PvRBP2b* (**Figure 2**). This observation raises intriguing questions regarding the potential fitness costs associated with hosting multiple copies of a specific gene within the parasite’s genome.

Our findings showed that individuals whose infections contained high copies of *PvDBP*1 likely experienced chills and shivering. Multiplications of *PvEBP/DBP2* were more related to muscle pain but less likely to joint pain, but higher copies of *PvRBP2b* were prone to joint pain and less likely to muscle pain. Increased incidence of joint pain with increased *PvRBP2b* copies may be explained by the high proportion of reticulocytes contained within the bone marrow and has been suggested as *P. vivax* reservoirs [48, 49]. While the exact cause of chills and shivering and muscle pain by *P. vivax* is unclear, these symptoms could be caused by an intense inflammatory reaction caused by pro-inflammatory cytokines released by the host during infection, although further research is needed to confirm why these differences may occur [50]. One of the inherent constraints encountered when working with clinical patient samples is the variability in the timing of data collection, typically coinciding with the patient’s presentation for treatment. This variability may arise from patients seeking treatment at different stages of infection, potentially before the complete manifestation of all associated symptoms.

To conclude, *PvDBP1*, *PvEBP/DBP2*, and *PvRBP2b* multiplications were common in Duffy-positive and Duffy-negative Ethiopians. Although Duffy-positive and Duffy-negative populations differ with the proportion of single- versus multi-copies, copy number differences were not correlated with parasitemia or gene expression. Further investigations on the functional significance and evolution of multiplications of these invasion genes across transmission settings will provide more insight into the epidemiological impact of copy number variations on *P. vivax* infections in Duffy-negative Africans.

## Ethics statement

The study protocol was reviewed and approved by the National Research and Ethics Review Committee (NRERC) (Ref. No.03/246/796/22). Written informed consent/assent was obtained from all consenting heads of household, parents/guardians, and individuals who were willing to participate in the study. All experimental procedures were performed following the IRB-approved protocol.

## Acknowledgements

We thank the field team from Jimma University for sample collection; the communities and hospitals for their willingness to participate in this research; and undergraduate students at UNC Charlotte who were involved in sample processing.

## Author Contributions

KP, JP, and EL conceived and designed the study. BA, DY, supervised data collection and manuscript review. KP, RR, JT, and EL collected and analyzed the data. KP, DK, and EL wrote and revised the paper. All authors read and approved the final manuscript.

## Table and Figure legends

**Table 1:** Frequency of single-copy and multi-copy between the Duffy-positive (T/T and T/C) and Duffy-negative (C/C) groups for *PvDBP1*, *PvEBP*/*DBP2*, and *PvRBP2b*.

**Table 2:** Odd’s Ratio for *PvDBP1*, *PvEBP/DBP2*, and *PvRBP2b* copy number variations in relation to clinical symptoms. *P*<0.01 are annotated with ***, *P*<0.05 are annotated with **, and *P*<0.1 are annotated with *.

**Supplementary Table 1**: List of primers, probes and plasmids which were used for *P. vivax* 18S SYBR, *DARC* Taqman, and *PvDBP1*, *PvEBP*/*DBP2*, and *RBP2b* CNV qPCR assays.

**Supplementary Table 2**: Compiled patient data including demographic data, qPCR raw data, and clinical symptoms data reported by patients.

**Supplementary Table 3**: Transcriptomic data for a subset of 10 *P. vivax* infected samples. Further analysis showed no significant relationship between copy number variations of *PvDBP1*, *PvEBP*/*DBP2*, and *PvRBP2b* gene expression levels.

