## Supplementary Table 1 for "Copy number variations of *Plasmodium vivax DBP1*, *EBP/DBP2*, and *RBP2b* in Duffy-positive and Duffy-negative Ethiopians"

| PCR | Primer Name | Sequence |
|---|---|---|
| *P. vivax* 18S | VIV_18S_F | 5’ – AGAATTTTCTCTTCGGAGTTTATTCTTAGATTGCT – 3’ |
| | FAL_VIV_18S_R | 5’ – GCCGCAAGCTCCACGCCTGGTGGTGC – 3’ |
| Duffy Genotyping Primers | DARC_Taq_F | 5’ – GGCCTGAGGCTTGTGCAGGCAG – 3’ |
| | DARC_Taq_R | 5’ – CATACTCACCCTGTGCAGACAG – 3’ |
| Duffy Genotyping Probes | DARC_C_FAM | FAM- CCTTGGCTCTTA**C**CTTGGAAGCACAGG-BHQ |
| | DARC_T_HEX | HEX- CCTTGGCTCTTA**T**CTTGGAAGCACAGG-BHQ |
| *PvDBP* Duplication | PvDBP_AF | 5’ – CCATAAAAGGTAGGAAATTGGAAA – 3’ |
| | PvDBP_AR | 5’ – GCATTTTATGAAAACGGTGCT – 3’ |
| | PvDBP_BF | 5’ – TCATCGAGCATGTTCCTTTG – 3’ |
| | PvDBP_BR | 5’ – TTGCACGTACTCGAAACTCAG – 3’ |
| | PvDBP_AF2 | 5’ – ACGCGATGTATCTTCTTTTCA – 3’ |
| | PvDBP_AR2 | 5’ – TAGAACGCACAGTTATTGGC – 3’ |
| Synthetic genes for CNV (For standard curve) | PvDBP | GAAAAACTGTAATTATAAGAGAAAACGTCGGGAAAGAGATTGGGACTGTAACACTAAGAAGGATG<br>TTTGTATACCAGATCGAAGATATCAATTATGTATGAAGGAACTTACGAATTTGGTAAATAATACA |
| | PvEBP/DBP2 | GGTGTAAAAATTGCGGGGGGAAGTACGAAAGATGAAAATCGGGGAGATCCAACATCGCACGAAAT<br>AAGATCGCATGAAGGAGGAAGTAGAGCAGCGGTCAATGGTCAAAGGGATGACGCTGGACGTGTTC |
| | PvRBP2b | ACACAAAGACCTCAATCCCATAAAGTCCGATATTGCAAATATAGAAACAGAATTGAAAAAGCATAGG<br>AAACTTTTCGAAATAGGACTTTTGAATAAGGTTATCGAGATAGCTAAAACGAGAAAATTGTACATGG<br>ATTCAGTGGAACAGCTCCTAAATTCCTCCATCAATAATTTTGTTACCCTCTTTAATGGGTTCAATTTAA<br>AAAAATATAAAATTGAAACCAGTTTAGACGTTTATAAAGGAAAAATAAGTGAATTCCACACTGAATT<br>CCTTCAGCCATACGAATCAATTCAAAATAAGGAAAAACGCGTACTGGACCCCTCGGTAGATTTTAGT<br>GCAGCAAAAAATATGCGAGAAGAAGCTCAAAAGGAGGAAAAAAATATCGAAAATGTAGAAAAAAA<br>GACAAAAAAATATCTAAGCGATATTAACAAAATGGAATCCTTTACATTCATACGTAATATGAGA |
| | β-Tubulin | CAGGAGTTACATGTTCGTTAAGATTTCCTGGTCAGTTAAATTCTGATTTGAGAAAATTAGCTGTCAAT<br>TTAATTCCCTTCCCAAGACTCCACTTTTTTATGATTGGTTTTGCACCACTAACAAGCAGAGG |
| Endogenous Control for CNV | CNV_β-Tubulin_F | 5’ – CATGTTCGTTAAGATTTCCTGGT – 3’ |
| | CNV_β-Tubulin_R | 5’ – GTTAGTGGTGCAAAACCAATCA – 3’ |
| *PvDBP* CNV | CNV_PvDBP_F | 5’ – AATTATAAGAGAAAACGTCGGGAAAG – 3’ |
| | CNV_PvDBP_R | 5’ – ACCAAATTCGTAAGTTCCTTCATACA – 3’ |
| *PvEBP/DBP2* CNV | CNV_PvEBP_F | 5’ – GGGGGAAGTACGAAAGATGAA – 3’ |
| | CNV_PvEBP_R | 5’ – CATCCCTTTGACCATTGACC – 3’ |
| *PvRBP2b* CNV | CNV_PvRBP2b_F | 5’ – GACCTCAATCCCATAAAGTCCG – 3’ |
| | CNV_PvRBP2b_R | 5’ – GGAATCCTTTACATTCATACG – 3’ |

**Supplemental Table 1**. Information of primers and plasmids for *P. vivax* screening, Duffy genotyping, and gene duplication assays.
